# Biological aging of different blood cell types

**DOI:** 10.1101/2024.05.07.592895

**Authors:** Saara Marttila, Sonja Rajić, Joanna Ciantar, Jonathan KL Mak, Ilkka S Junttila, Laura Kummola, Sara Hägg, Emma Raitoharju, Laura Kananen

## Abstract

A biological age (BA) indicator is intended to capture detrimental age-related changes occurring with passing time. To date, the best-known and used BA indicators include DNA-methylation-based epigenetic ages (epigenetic clocks) and telomere length. The most common biological sample material for epidemiological aging studies is composed of different cell types, whole blood. We aimed to compare differences in BAs between blood cell types and assessed BA indicators’ cell type-specific associations with donor’s calendar age.

Analysis on DNA methylation-based BA indicators including telomere length, methylation level at cg16867657 (a CpG-site in *ELOVL2*) and the Hannum, Horvath, DNAmPhenoAge and DunedinPACE epigenetic clocks was performed in 428 biological samples from 12 blood cell types. BA values were different (p<0.05) in the majority of pairwise comparisons between the cell types. Most cell types also displayed differences as compared to whole blood (p<0.05). Some of the observed differences persisted across blood donor’s calendar ages from 20 to 80 years (50-years-difference in DNAmPhenoAge between naïve CD4+ T cells and monocytes), while others did not (up to four-fold difference in DunedinPACE values between monocytes and B cells). All BA indicators, except DunedinPACE, had mostly a very strong correlation with donor’s calendar age within a cell type.

Our findings demonstrate that DNA methylation-based indicators of biological age exhibit cell type-specific characteristics, underscoring the importance of accounting for cell composition in related studies. Our results have implications for understanding the molecular mechanisms underlying epigenetic clocks and and provide guidance for utilizing them as indicators for success of aging interventions.

## Introduction

By definition, biological age (BA), or an aging biomarker, should better predict future health status as compared to chronological age. By AFAR (American Federation for Aging Research) criteria “It must monitor a basic process that underlies the aging process, not the effects of disease.”^1–4^. Of the many established BA indicators ^4,5^, the most well-known and used are DNA-methylation-based epigenetic ages (epigenetic clocks) and telomere length, a hallmark of aging^6^. Ideally, these indicators should reflect how health interventions influence biological aging. However, the underlying molecular mechanisms of the epigenetic clocks are still unknown.

Accelerated biological aging (or aging rate) indicated by telomere length^7^ and epigenetic clocks^6^ predicts health span, lifespan or both in large-scale cohort studies. Typically, these analyses are performed in whole blood samples that are mixtures of various blood cell subtypes. As such, blood cell composition is a potential confounder in the analyses, because blood cell composition changes with advancing age ^8^, already before middle-age ^9^. Typical characteristics of age-related remodeling of the immune system include decreasing naïve CD8 and CD4 T cell and increasing exhausted CD28-T cell counts, declining CD4 to CD8 T cell ratio, and potentially also altered NK cell count and functionality ^10,11^. This remodeling is highlighted by the existence of IMM-AGE^12^, a blood cell composition-based, potential BA indicator. Further, changes in blood cell composition are seen in many age-related conditions (e.g. frailty ^13^) and diseases *e.g.* cancer ^14^, Alzheimer’s ^15^, and cardiovascular diseases ^16,17^. The age-related remodeling of the blood cell composition is not limited to these changes, but these are the well-known examples for which there is epidemiological evidence of their relationship to aging and aging phenotypes.

A better understanding on the biological aging at the cell subtype-level within tissues is needed. Previous studies have shown telomere length ^18–21^ and DNA methylation level at cg16867657, a CpG site in *ELOVL2*^22^, are tissue- and cell type-specific in their absolute values and age-related changes. A few previous studies have shown that epigenetic ages by DNAmPhenoAge and Horvath differ between blood cell types^23^, but BA or biological aging rate indicated by epigenetic clocks developed more recently are studied less in separated cell subtypes. Importantly, previous such analyses have been made typically using separated cells originating from different individuals and datasets with less than 10 individuals each^24^. Further, it is unknown in what way cell subtype-specific epigenetic age values indicated by the 2^nd^ and 3^rd^ generation epigenetic clocks change with advancing calendar age across adulthood. Thus, in this study, we aimed to **1)** assess differences in values of DNA-methylation-based BA indicators between blood cell types originating from the same blood donors and with more adequate sample size. **2)** We also aimed to assess BA indicators’ cell type-specific associations with donor’s calendar age. The BA indicators included ‘the 1^st^ generation clocks’ (ELOVL2-CpG-site, cg16867657 ^23^, Horvath ^26^ and Hannum ^27^), ‘the 2^nd^ generation clock’ (DNAmPhenoAge^28^), ‘the 3^rd^ generation clock’ (DunedinPACE ^29^) as well as telomere length (DNAmTL, estimated based on DNA methylation data ^30^). In our main analyses, we performed pairwise comparisons for BA indicator values between whole blood, peripheral blood mononuclear cells (PBMCs) and up to ten separated blood cell subtypes in four separate data sets with 428 biological samples, originating from the same blood donors. Then, we assessed cell subtype-specific associations of the different BA indicators with calendar age. In our additional analyses, we repeated pairwise comparison analysis with principal component derivates of the clocks^31^, assessed cell subtype-specific correlations of the different BA indicators with each other, and last, exemplified blood cell subtype count trajectories over decades in a longitudinal cohort sample (The Swedish Adoption/Twin Study of Aging [SATSA], n=328).

## Methods

### Data sets

We included four datasets available in NCBI GEO ^32,33^ (GSE35069 ^34^, GSE131989 ^35^, GSE166844 ^23^, GSE78942 ^36^) in which DNA methylation data was available from separated immune cell subtypes (Table 1) for this study. These subtypes were separated using fluorescence-activated cell sorting (FACS) as described in detail in the original publications ^23,34–36^. Surface markers used for the FACS analyses are summarized in Supplementary Table S1. We included only datasets in which the different immune cell populations were available from the same individuals as complete cases. For the cell count trajectory analysis, DNA-methylation-based cell count estimates in whole blood samples in the Swedish Adoption/Twin Study of Aging (SATSA, n=328 with 657 observations, baseline ages 48-98, mean age 68.5) were used ^37^.

**Table 1.** Data sets in pairwise comparisons. Details on how different cell types were separated are provided in Supplementary Table 1.

| **Data set** | **Individuals,**<br>**n** | **Calendar age**<br>**(years)** | **Calendar age**<br>**available for**<br>**each individual** | **Female,**<br>**%** | **Cell**<br>**sample**<br>**types,**<br>**n** | **Available cell sample types** | **Pairwise**<br>**cell type**<br>**comparisons,**<br>**n** |
| --- | --- | --- | --- | --- | --- | --- | --- |
| **GSE131989** | 49 | 54 ± 17.5 | Yes | 100 | 4 | CD14+, CD19+, CD4 memory T cells, CD4 naïve T cells | 6 |
| **GSE166844** | 28 | 19 ± 0 | Yes | 40 | 6 | Whole blood, Granulocytes, CD14+ monocytes, CD19+ B cells, CD8+ T cells, CD4+ T cells | 15 |
| **GSE35069** | 6 | 38 ± 13.6 | No | 0 | 10 | Whole blood, PBMC, Granulocytes, Neutrophils, Eosinophils, CD14+ monocytes, CD19+ B cells, CD56+ NK cells, CD8+ T cells, CD4+ T cells | 45 |
| **GSE78942** | 24* | 62.1 ± 9.9 | No | NA | 2 | CD4+CD28+ T cells, CD4+CD28- T cells | 1 |

\*DNA methylation measured from two pooled samples of separated cells; 12 individuals in a sample

### BA indicators

We assessed different BA indicators using DNA methylation data from the aforementioned datasets (Table 1). The indicators of BA (or biological aging rate) investigated were telomere length estimated based on DNA methylation (DNAmTL) ^30^, methylation level of *ELOVL2* at one CpG (cg16867657)^25^, Hannum^27^, Horvath^26^, DNAmPhenoAge^28^ and DunedinPACE^29^ as well as the principal component derivates of Horvath, Hannum, DNAmPhenoAge and DNAmTL^31^. In three of the included datasets, DNA methylation was measured using Illumina 450K (GSE35069, GSE13198) or Illumina EPIC (GSE166844) array, allowing us to calculate all the ten indicators of BA. In dataset GSE78942, methylation data were measured using Illumina 27K array, allowing us to calculate only Horvath and DNAmPhenoAge. All BA indicators were calculated from the normalized and preprocessed data available in GEO.

Horvath (for datasets GSE35069, GSE13198 and GSE166844), Hannum, DNAmPhenoAge and DNAmTL (for all included datasets) were calculated using the DNAmAge function of the methylclock package version 0.8.2^38^. For GSE78942, Horvath was calculated using the webpage tool available in https://dnamage.clockfoundation.org/. DunedinPACE was calculated as described in the original publication^29^ with the R package DunedinPACE. The principal component derivates of the clocks were calculated as previously described ^31^. Methylation value of the probe cg16867657 in *ELOVL2* was extracted directly from methylation data available in GEO for each dataset.

### Statistical analysis

Statistical significance for the pairwise comparisons were assessed using the Mann Whitney U-test. BA values were compared between the cell subtypes at group-level within a data set. Cell subtype-specific BA values were visualized as boxplots with dots and line plots, and pairwise differences as boxplots. Cell subtype-specific relationships between values of different BA indicators and calendar age were assessed using correlation statistics (Spearman), and the relationships were visualized as scatterplots.

In our additional analyses, we assessed cell subtype-specific relationships between values of different BA indicators using correlation statistics (Spearman) and the relationships were visualized as scatterplots. In the longitudinal cohort data, cell subtype count trajectories were visualized as line plots and significance for the cell count change with calendar age was obtained using mixed linear model. In GSE131989 and SATSA, calendar age was used as individual-level phenotypic data in our statistical analyses. Data were analysed and visualized using R statistical software (version 4.2.2) and R-packages ggplot2. P-value threshold for statistical significance was set to 0.05.

## Results

### Pairwise comparisons

BA values for each cell type in the different data sets (Table 1) in our analysis are shown in Figure 1, Table 2, Supplementary Figure S1 and Supplementary Table S2. We performed as our main analysis pairwise comparisons of the BA indicator values between the blood cell subtypes. In summary, BA values, including principal component derivates of the epigenetic clocks, were different (Mann-Whitney U-test p<0.05) in the majority of pairwise comparisons between the cell types (Table 2, Figure 2, Supplementary Table S3-S5, Supplementary Results). Most cell types also displayed differences as compared to whole blood (Mann-Whitney U-test p<0.05, Figure 2, Supplementary Table S3-S5). Some of the observed differences persisted across blood donor’s calendar ages from 20 to 80 years, for example the 50-years-difference in DNAmPhenoAge values between naïve CD4+ T cells and monocytes (Figure 3). However, for example, the up to four-fold difference in DunedinPACE values between monocytes and B cells did not persist over time (Figure 3).

**Figure 1.**
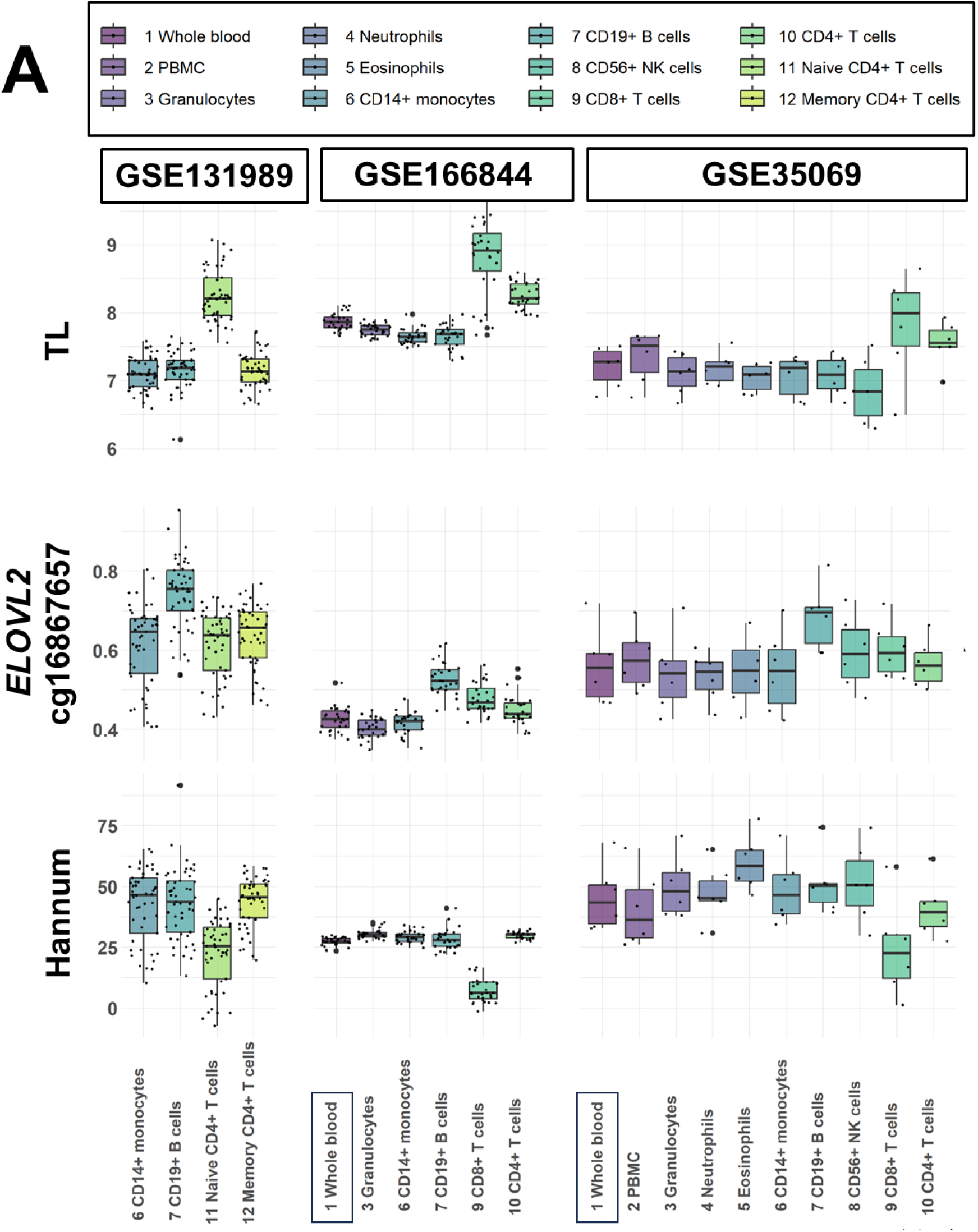
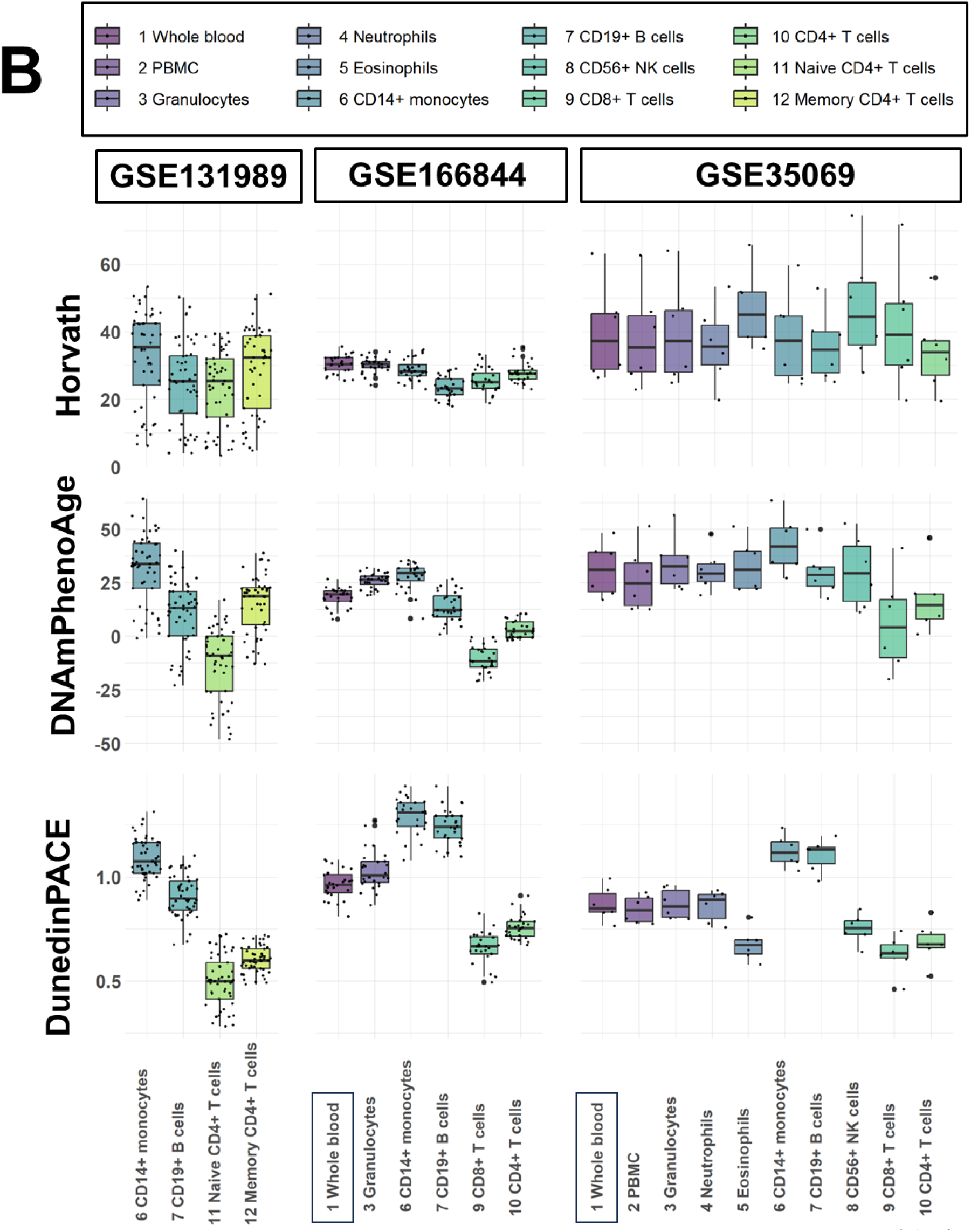
Blood cell type-specific biological ages (BAs) and a BA rate. Values of DNAmTL (TL), cg16867657 in *ELOVL2*, and Hannum are summarized as boxplot with dots in panel A, and Horvath, DNAmPhenoAge and DunedinPACE in panel B. These DNA methylation-based BA indicators were assessed in three DNA methylation data sets (GSE131989, GSE166844, GSE35069) with 424 biological samples from 83 individuals and including 12 cell sample types. Boxes are colored according to cell type (1-12). Each dot represents one individual.

**Figure 2.**
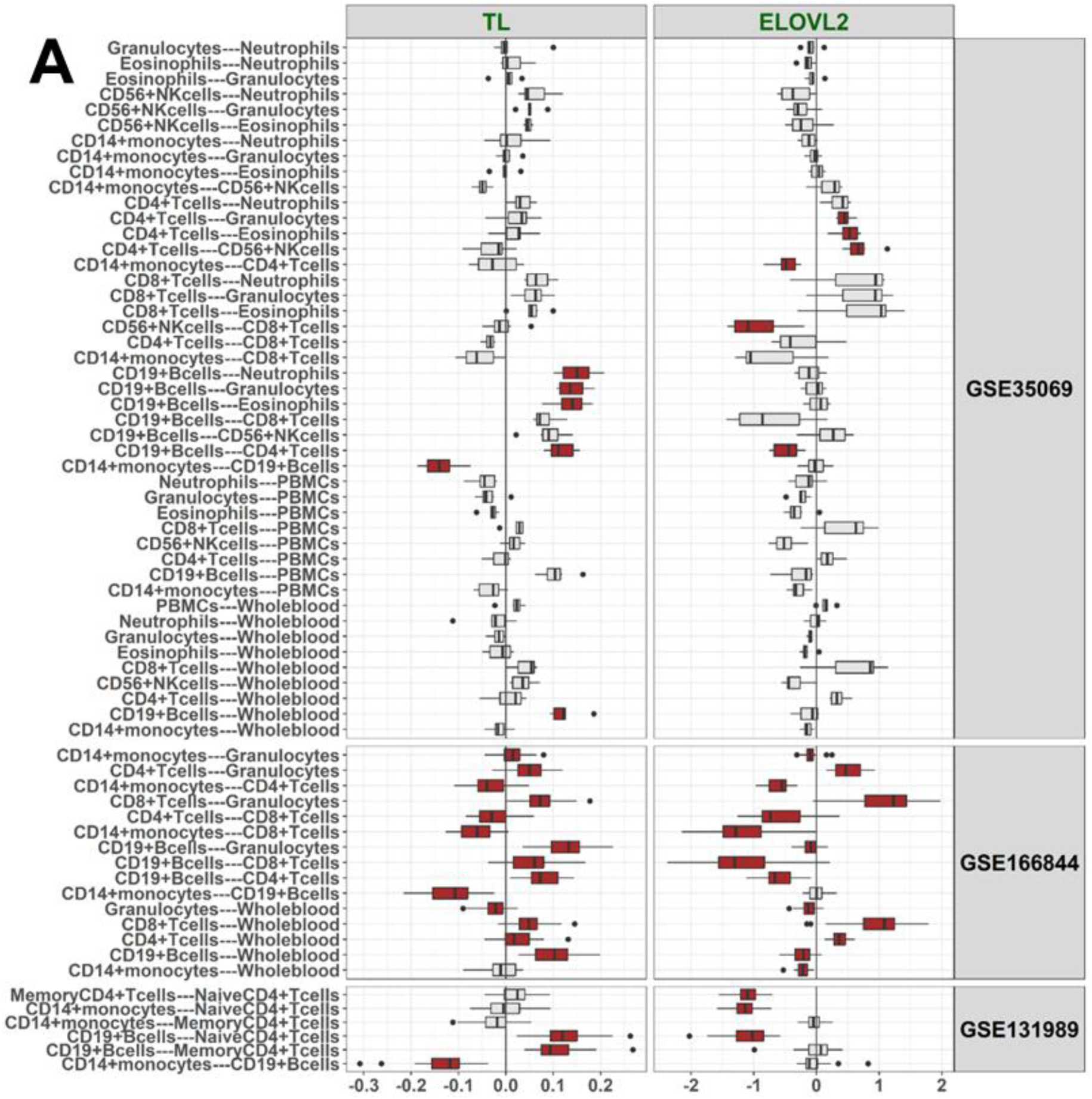
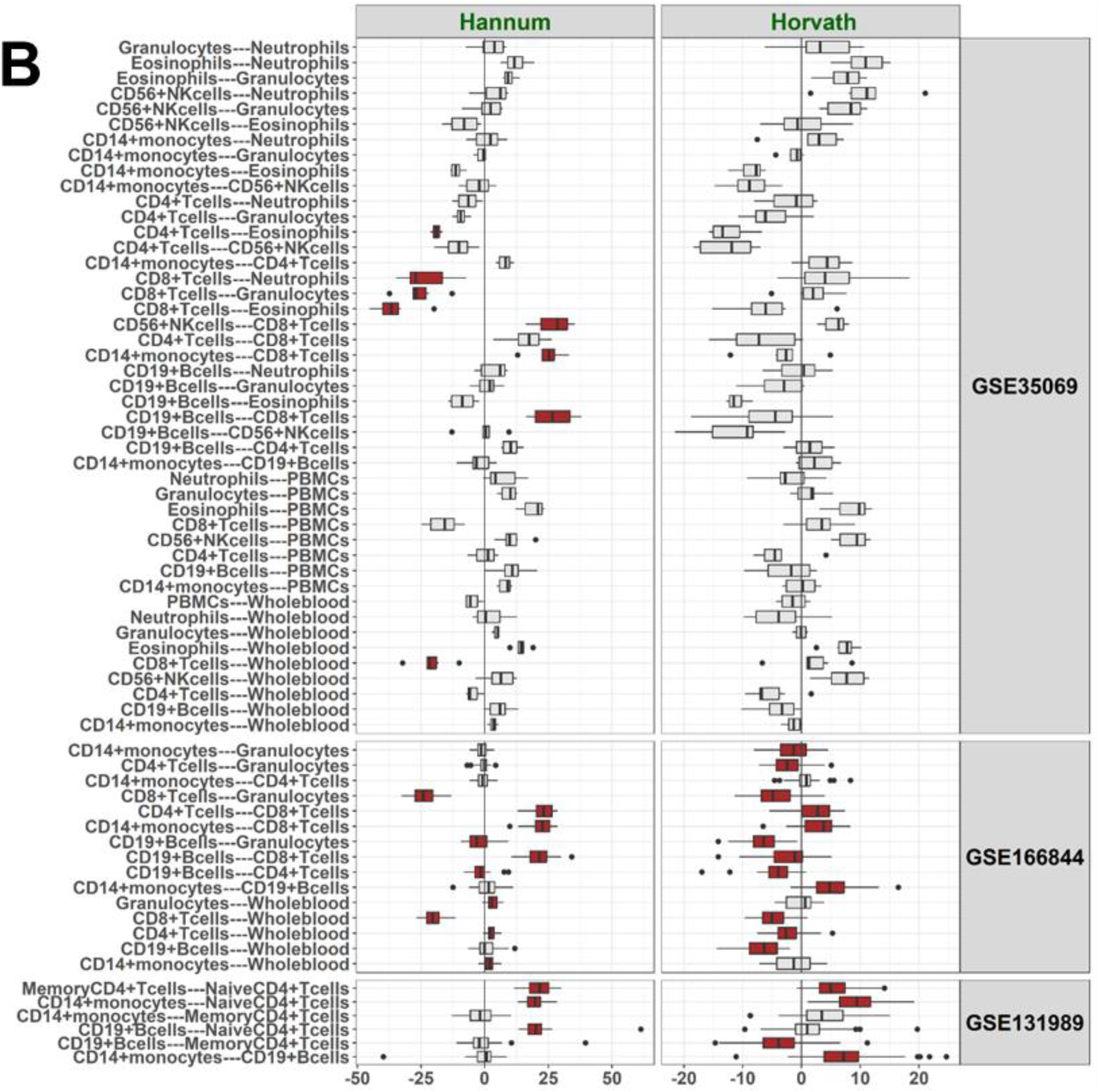
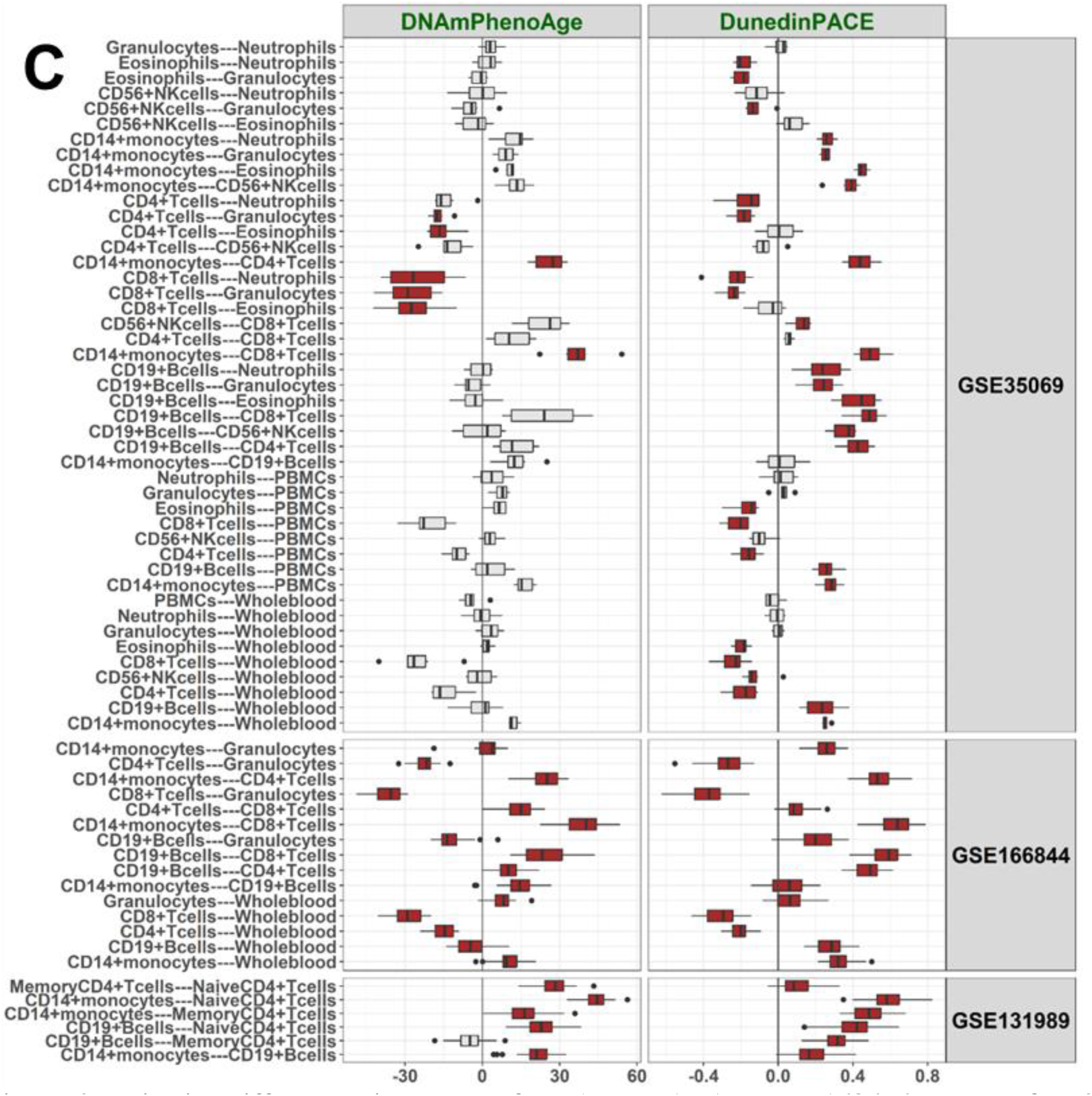
Pairwise differences in values of DNAmTL (TL) and cg16867657 at *ELOVL2* (ELOVL2) (A), Horvath and Hannum (B), and DunedinPACE and DNAmPhenoAge (C) between the cell types. Cell pairs with statistically significant difference in BA values (Mann Whitney U, p<0.05) are colored with red, otherwise with grey. Difference in BA indicator values between a cell-pair (Δ) was calculated for each individual and these differences are shown as boxplots for the three data sets. Δ-value for a BA indicator is shown on the x-axis. In GSE131989 including 49 blood donors, there were 6 cell type pairs, in GSE166844 including 28 blood donors, 15 pairs, and in GSE35069 including six blood donors, 45 cell type pairs to be compared. Cell type-specific BA values within a dataset are summarised in Figure 1, Supplementary Figure 1 and Table S2, and p-values for the comparisons are presented in Supplementary Tables S3-S5.

**Figure 3.**
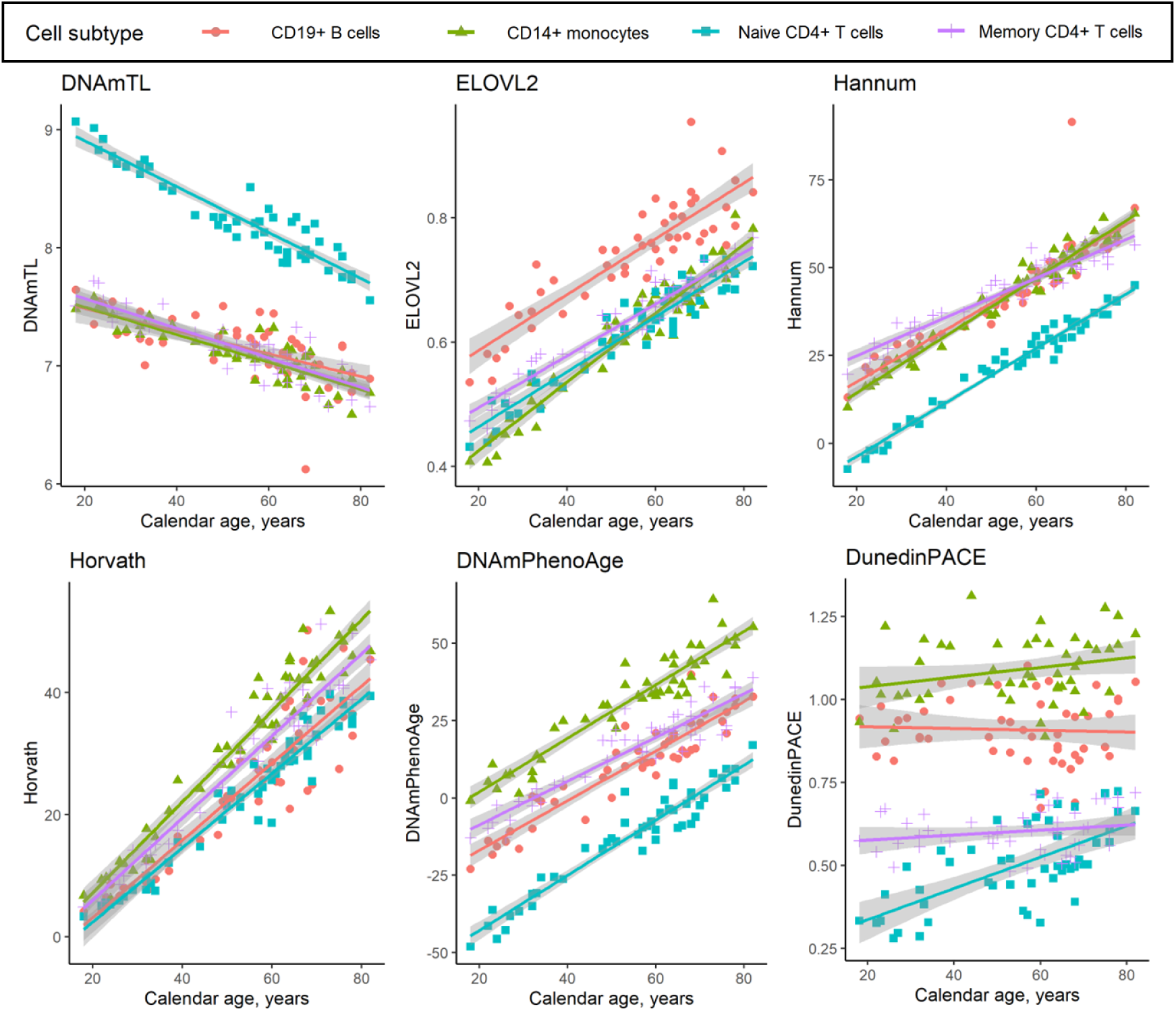
Cell type-specific associations of DNA-methylation-based BA indicators (and biological aging rate) with blood donors’ calendar age in GSE131989. DNA methylation data within four separated cell subtypes (CD19+ B cells, CD14+ monocytes, naïve CD4+ T cells, memory CD4+ T cells) originated from 49 individuals aged 18-82 years (all females). Correlation statistics showing the significance for the associations with calendar age are shown in Supplementary Table S6. Grey areas indicate 95% Confidence Intervals for the linear fit lines.

**Table 2.** BA values according Horvath and DNAmPhenoAge in displayed ‘younger’ values in CD4+CD28+ T cells as compared to CD4+CD28-T cells in GSE78942. DNA methylation was measured from four pooled biological samples of purified cells. In pool1, cells were separated from 12 individuals and the cells were pooled as two biological samples (CD28+ and CD28-cells), and in pool2, cells originated from other set of 12 individuals and the separated cells were pooled in a similar way as pool 1. Calendar age of these healthy blood donors was 45-75 years (mean[SD]=62.1[9.9]).

| Dataset | Sample type, sample id | Horvath | DNAmPhenoAge |
| --- | --- | --- | --- |
| GSE78942 | CD4+CD28- T cells, pool1 | 65.9 | 50.8 |
|  | CD4+CD28- T cells, pool2 | 72.9 | 57.0 |
|  | CD4+CD28+ T cells, pool1 | 39.0 | 8.5 |
|  | CD4+CD28+ T cells, pool2 | 35.1 | 10.7 |

As expected, pairwise comparisons were more often statistically significant (Mann-Whitney U test p-value <0.05) in GSE166844 and GSE131989 with larger number of individuals than in GSE35069 which included six individuals (Table 1, Table 3). Other important details for results interpretation are that GSE131989 and GSE35069 included individuals with wide calendar age range while individuals in GSE166844 were all 19 years old. While most cell types were available in at least two datasets, neutrophils, eosinophils and CD56+ NK cells were available for analysis in GSE35069 only. In GSE78942, the difference in BA values was apparent but statistical analysis was not possible as it comprised four biological samples only.

**Table 3.** Number of cell pairs showing differences for each indicator of BA in three datasets. Mann-Whitney U test p-values are shown in Supplementary Table S3-S5.

|  | Data set |  |  |
| --- | --- | --- | --- |
|  | GSE131989 | GSE166844 | GSE35069 |
| Number of pairwise comparisons | 6 | 15 | 45 |
| Number of cell pairs showing pairwise differences(%) |  |  |  |
| DNAmTL | 3(50) | 14(93) | 6(13) |
| ELOVL2 | 3(50) | 14(93) | 6(13) |
| Hannum | 3(50) | 10(67) | 8(18) |
| Horvath | 4(67) | 12(80) | 0(0) |
| DNAmPhenoAge | 5(83) | 15(100) | 7(16) |
| DunedinPACE | 6(100) | 15(100) | 31(69) |

### CD19+ B cells

Methylation data in CD19+ B cells was available in three datasets. As compared to other cell types, CD19+ B cells displayed a statistically significant difference (Mann-Whitney U test p-value <0.05) in majority of the pairwise comparisons in GSE131989 and GSE166844 (Figure 2, Supplementary S3-S4). In the smallest dataset, GSE35069, statistically significant differences were mainly observed for DunedinPACE (Figure 2, Supplementary Figure S1, Supplementary Table S5). In summary, our results suggest CD19+ B cells are, according to the studied BA indicators, ‘younger’ as compared to CD14+ cells, but ‘older’ as compared to naïve CD4+ cells and total CD8+ T cells, although there are some discrepancies between the different BAs (Figure 2, Supplementary Table S3-S5). In comparison to whole blood, no clear pattern was observed for CD19+ B cells.

### T cell subsets

Data on various subsets of T cells was available in four datasets, including total CD4+ and CD8+ T cells (GSE166844 and GSE35069), CD4+ naïve and memory T cells (GSE131989) and CD4+CD28- and CD4+CD28+ T cells (GSE78942). Majority of pairwise comparisons across these cell types were statistically significant (Figure 2, Table 2, Supplementary Tables S3-S5). Our results suggest that CD8+ T cells are ‘younger’ as compared to CD4+ T cells, and that naïve CD4+ T cells are ‘younger’ as compared to memory CD4+ T cells (Figure 2, Supplementary Table S3-S5). In addition, CD4+CD28+ cells were identified to be ‘younger’ as compared to CD4+CD28-according to both BA indicators available for this dataset, Horvath and DNAmPhenoAge (Table 2), although for this data no statistical tests could be performed, as there were only four biological samples. As compared to whole blood, both CD4+ and CD8+ T cells are ‘younger’, although there are discrepancies between different BA indicators (Figure 2, Supplementary Table S3-S5). The magnitude of difference was larger between CD8+ T cells and whole blood as compared to CD4+ T cells and whole blood (Figure 1, Figure 2).

### CD14+ monocytes

Data on CD14+ monocytes was available in three datasets. As compared to other cell types, majority of pairwise comparisons between CD14+ monocytes were statistically significant in GSE166844 and GSE131989 (Figure 2, Supplementary Figure S1, Supplementary Table S3 and S4). Our results suggest that CD14+ monocytes are ‘older’ as compared to various T cell subsets, ‘older’ as compared to CD19+ B cells, and also ‘older’ as compared to whole blood samples (Figure 2, Supplementary Tables S3-S5).

### Cell-subtype specific BA values across adult calendar ages

All BA indicators, except DunedinPACE, correlated strongly with calendar age within a cell type in dataset GSE131988 (>0.8 or < -0.7, Figure 3, Supplementary Table S6). DunedinPACE values increased most consistently with a higher calendar age within naïve CD4+ T cells (Spearman’s ρ=0.636), but in other cell types tested the correlations were more modest or nonsignificant (Figure 3, Supplementary Table S6). The analysis could only be performed in this one dataset, as calendar age was not available, or all individuals were of the same age, in others.

### Additional analyses

#### Pairwise comparisons for principal component clocks

The epigenetic clocks have been reported to suffer from technical noise^31^. The proposed solution is to utilize principal components instead of the individual level CpG data to calculate the clocks i.e. PC-clocks. To verify that the observed differences in BA indicators across cell types are not due to technical noise of the Illumina array, as an additional analysis, we repeated the pairwise comparison analysis with the principal component derivates for Horvath, Hannum, DNAmPhenoAge and DNAmTL (Supplementary Table S2). Our results show that the observed differences between the cell types in the main analysis remained significant for the studied PC-clocks (Supplementary Table S3, S4 and S5).

#### Relationships between different BAs

Then, we explored relationships between the values of different BA indicators (Supplementary Figure S2) and focused on the relationships within each cell subtype population (Supplementary Table S6-S8). The majority of BA indicators showed strong or very strong correlations (>0.7 or < -0.7) with each other within the different cell subtype populations in GSE131989 and GSE35069 (Supplementary Table S6 and S8), which have wide age ranges. However, only a very few moderate or stronger correlations (>0.5 or < -0.5) were observed in GSE166844 (Supplementary Table S7), which includes individuals with the same calendar age. An exception in the cell type-specific correlations was seen for DunedinPACE as the correlations were, overall, lower or non-existing (Figure 3A, Supplementary Table S6-S8).

#### Blood cell composition trajectories

In the last additional analysis, we visualized estimated blood cell composition trajectories in a longitudinal cohort (SATSA) with decades of follow-up (Supplementary Figure S3) and observed changes in cell counts with advancing calendar age for all blood cell subtypes that were in our pairwise comparisons and also available in SATSA (p<0.005). The counts of B cells, CD4+ and CD8+ T cells, naïve CD4+ and CD8+ T cells decrease, while the counts of CD8+CD28-CD45RA-T and NK cells, plasmablasts, monocytes and granulocytes increase from midlife into old age (Supplementary Figure S3).

## Discussion

We assessed ten DNA-methylation-based BA indicators, Horvath^26^, Hannum^27^, DNAmPhenoAge^28^, DNAmTL^30^, their principal component derivates ^31^, DunedinPACE^29^ and methylation level of *ELOVL2* at cg16867657^25^ in 428 biological samples, in up to 12 blood cell types, collected and separated from the same set of individuals. Our results show a significant difference (p < 0.05) in BA values, including principal component derivates of the epigenetic clocks, in the majority of pairwise comparisons between the cell types and as compared to whole blood. As a new finding, we show that the cell type-specific BA values of the blood cells appear to persist across human adulthood, with the exception of DunedinPACE. For example, the 50-years-difference in DNAmPhenoAge values between naïve CD4+ T cells and CD14+ monocytes, persists across calendar ages from 20 to 80 years. To put the 50-years-difference into perspective, the BA value difference is approximately 60 years between a 20- and 80-years-old person, but the cell type-specific difference is a few years between two persons with the same calendar age. Thus, in line with Zhang et al (2023)^24^, we conclude that calendar age and blood cell composition together explain the great majority of variation in BA values. As an exception among the BA indicators, the DunedinPACE values can have up to four-fold differences between the cell types, but the differences appear not to persist across human life course across all cell types studied here. Furthermore, by using longitudinal cohort data, we highlight how thoroughly blood cell composition changes with age during adulthood, in line with previous reports ^12,39–46^. The synthesis of this research evidence implies that the proportion of many of the cell types with ‘younger’ BA values in blood circulation, such as naïve CD4+ and naïve CD8+ T cells, decline with advancing calendar age, while the proportion of cells with mostly ‘older’ BA values, such as monocytes, become more prevalent (Figure 4).

**Figure 4.**
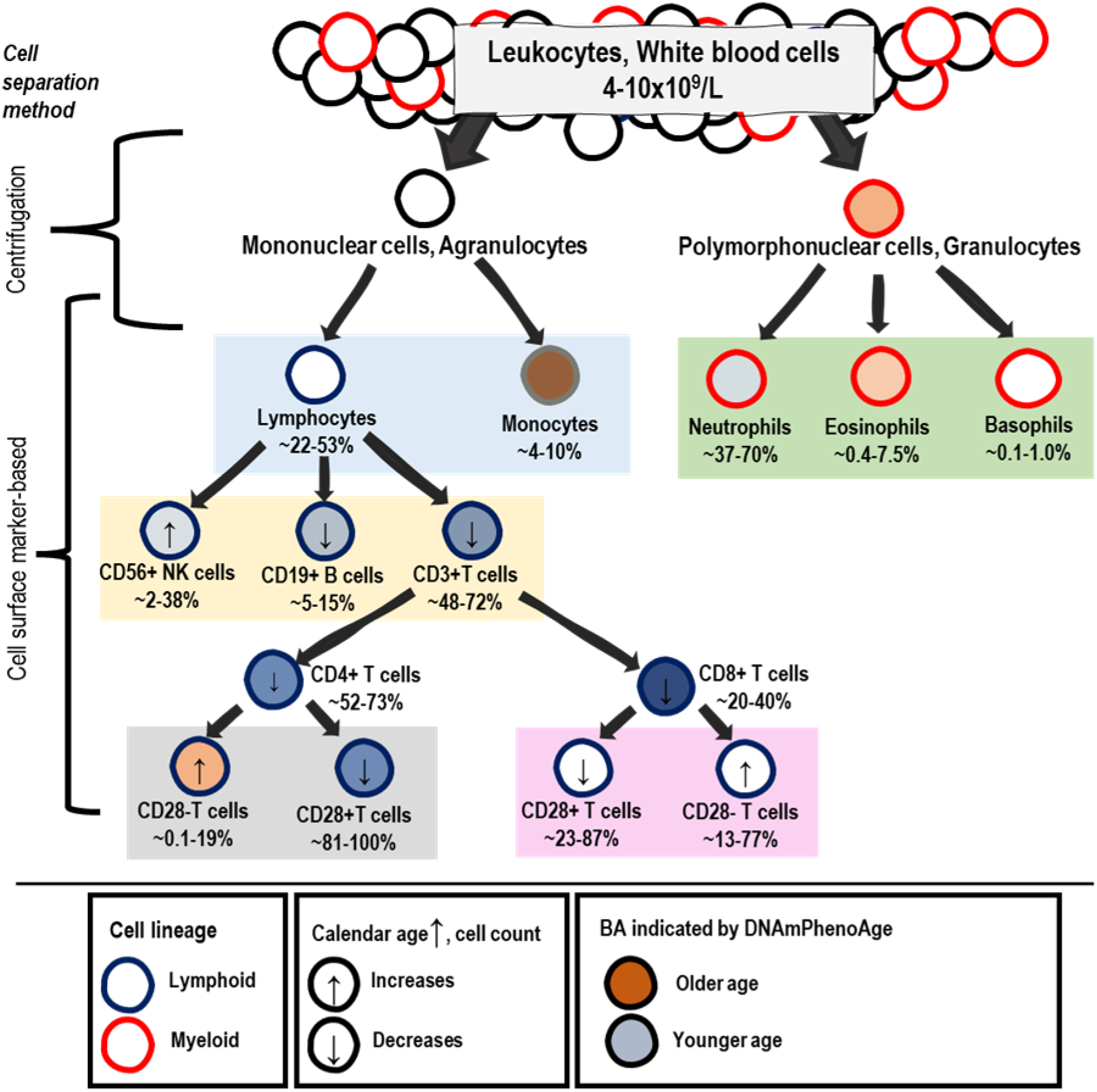
Graphical summary of typical blood cell subtype separation with cell type-specific biological ages (BAs), as well as cell proportion ranges and age-related changes at human population-level. Cell count prevalence ranges and changes with advancing age at population-level are according to previous reports^39,40,42,44–47^ and Supplementary Figure S3. Biological age (BA) indicated by DNAmPhenoAge is colored according to rank-orders of cell type-specific group mean values for DNAmPhenoAge in GSE131989, GSE166844, GSE35069 and GSE78942 in this study (Table 2, Supplementary Table S2). Our results suggest a trend of cell types with ‘older’ BA values increasing in numbers with increasing calendar age, and vice versa, cell types with ‘younger’ BA values decreasing with age.

So far, reports on blood cell type-specificity in Horvath, DNAmPhenoAge and DunedinPACE values have been based on pairwise comparisons between cell subtypes originating from different individuals^24^, small datasets (number of individuals<10)^24^, on whole blood data, where cell proportions have been estimated using deconvolution methods^24,48^ or for a single BA indicator at a time^23,43^. The strength of our approach was the inclusion of purified cell populations from four independent data sets, six BA indicators and the PC-clocks, and datasets consisting of the same sets of individuals for each cell type. Furthermore, we were able to assess relationships between DNAmPhenoAge and DunedinPACE values and donor’s calendar age within a cell type for a larger number of individuals than in previous studies.

Our observations are in line with previous studies^23,24,43,48^, in those parts where comparable. In our analysis, subsets of T cells, especially naïve CD4+ and total CD8+ T cells displayed generally the ‘youngest’ values of different BA indicators. Both CD19+ B cells and CD14+ monocytes displayed ‘older’ BA indicator values as compared to T cells, and of the two, CD14+ monocytes displayed the ‘oldest’ BA values. The differences between CD4+CD28+ and CD4+CD28-T cells were especially pronounced. That is, naïve CD4+ T cells showed ‘younger’ BA as compared to memory CD4+ T cells, and that CD4+CD28+ T cells showed ‘younger’ values of Horvath and DNAmPhenoAge as compared to CD4+CD28-T cells, and the differences were up to 40 years.

We identified statistically significant differences for the Horvath pan-tissue clock^26^ in the majority of pairwise comparisons in two independent data sets in line with our previous findings^43^ as well as other literature^23,24,48^. This finding is interesting as this 1^st^ generation clock was trained with data from altogether 53 somatic tissues^26^, and one could expect that different cell types would display similar values of this BA. In Kananen et al. (2016), Horvath values were higher with a higher FACS-analysis-based proportions of CD4+CD28-T cells as compared to CD4+CD28+ T cells when assessed from cells originating from individuals with the same calendar age. The other previous studies have reported up to twenty-years difference in Horvath values between different cell subtypes ^23,24,48^.

Zhang et al. (2023) reported lowest values for Horvath, Hannum, DNAmPhenoAge and DunedinPACE for naïve CD8+ T cells. Our datasets did not include naïve CD8+ T cells, only total CD8+ T cells, and they were generally observed to have lower values of BAs as compared to whole blood. In addition, in both datasets containing CD8+ T cells, they showed the lowest values of BAs among the different cell types. It is important to note that although naïve T cells are more prevalent in blood than CD28-T cells, especially in younger calendar ages ^41,49^, and we observed dramatic differences in their BA values when compared to whole blood, while BA values of CD4+CD28-T cells are closer those of whole blood. Thus, the magnitude of the possible contribution by naïve T cells to the BA values in a whole blood sample is substantial.

The DunedinPACE^29^ values, when measured in whole blood, have been shown to increase with a higher calendar age, even though this association is much weaker as compared to other epigenetic clocks^50^. The association between the donor’s calendar age and DunedinPACE^29^ values within a separated blood cell type has been assessed previously in data with only a few individuals^24^. Our correlation statistics from dataset (GSE131989) with 49 blood donors indicate that the association between DunedinPACE and calendar age may not be the same in all cell types. In naïve CD4+ T cells, the Spearman’s correlation ρ was 0.64 while in memory CD4+ T cells, CD14+ monocytes and CD19+ B cells, the correlation was weak or non-existing (Spearman’s ρ<0.3). This highlights the need to further study the effect of naïve CD4+ T cell counts on the DunedinPACE values measured in whole blood samples. For the other BA indicators (DNAmTL, methylation level at the *ELOVL2* CpG-site, Hannum, Horvath and DNAmPhenoAge), cell type-specific values correlated with calendar age strongly or very strongly (Spearman’s ρ >0.7 or < -0.7).

The majority of our results show a similar direction and magnitude for pairwise comparisons in the different BA values between the cell types. Certain cell types were either ‘younger’ or ‘older’ according to most of the indicators, for example, naïve CD4+ T cells were very often ‘younger’ than the other cells. The similarities might be explained by the fact that these BA indicators are based on DNA methylation which is tightly linked with cellular identity^51^. In parallel, according to some BA indicators, such as DNAmPhenoAge, monocytes are ‘older’ than B cells, naïve and memory CD4+ T cells, but according to Hannum they are not. The differences for the BA indicators may be explained by the fact that the different BAs are representing different domains in biological aging (e.g. DNA methylation in a gene vs telomere length vs epigenetic clocks) and of course, utilize varying sets of DNA methylation sites in the genome. Further, the epigenetic clocks can also be categorized into generations depending on the building strategy. The 1^st^ generation epigenetic clocks, such as Horvath ^26^ and Hannum clocks ^27^ were built to predict calendar age, the 2^nd^ generation epigenetic clocks, such as DNAmPhenoAge^28^, were built to predict biological age utilizing biomarkers and calendar age, while the 3^rd^ generation clock, DunedinPACE ^29^ was built to predict pace of aging, utilizing longitudinal biomarker and health data, and not calendar age as such. Horvath was trained in blood and multiple tissues^26^, and the rest are only based on measurements from blood samples.

The significance of cell proportion for epigenetic ages has been noted, to some extent, in previous literature and is an important consideration for the concepts of intrinsic and extrinsic epigenetic ages^52^. These measures of aging are both residual values of an epigenetic clock, such as Horvath or Hannum, after adjusting for calendar age, but intrinsic epigenetic age aims to be independent of blood cell composition as the composition is adjusted for. However, for the extrinsic epigenetic age, the cell composition is incorporated into its values as an additive element. Thus, extrinsic age is not intended to be a measure of the deep cellular mechanism in the aging process, but it is a composite measure. In a meta-analysis of 13 cohorts by Chen et al. (2016), extrinsic age values resulted a higher hazard ratio for mortality with more narrow confidence intervals than intrinsic age^52^. This implies that cell counts may give additive value for, for example, lifespan prediction, and the cell composition is not solely a potential confounding factor.

DNA methylation-based BA indicators are often developed for and measured in whole blood or PBMC samples. They can used in trials or interventions targeted at rejuvenation or reversing biological aging, but they can also be used to study physiological or pathological conditions not related directly to ageing as such. As ageing and various other physiological or pathological conditions can have an effect on blood cell composition, great care should be taken to disentangle the relationship between cell composition and these indicators. For example, a physically active lifestyle has been reported to rejuvenate the immune system by increasing the numbers of naïve T lymphocytes or by altering the CD4/CD8 ratio^53^. Fahy et al. (2019) have reported reversal of epigenetic aging in PBMCs indicated by four different epigenetic clocks with a thymus regenerating treatment. In a parallel analysis, they showed that treatment-related changes in circulating blood cell types include a decrease in monocytes and an increase in naïve CD4+ and CD8+ T cell, but did not account for the cell counts in the statistical analysis for the epigenetic clocks^54^. As our results indicate that monocytes have ‘older’ BA values while naïve T cells have ‘younger’ values, their results on the epigenetic clocks may have been influenced by the changes in immune cell proportions. In other studies on potential aging interventions, cell proportions have not been taken into account ^55^ or only the baseline cell proportions have been accounted for^56^. In general, when interpreting the results of potential aging interventions, great care should be taken to define what is meant and aimed by rejuvenation. Is the aim to change the cells’ intrinsic processes or not? One can ask, is a change in immune cell proportions alone a sufficient outcome for an intervention to be considered successful?

As an example of a physiological condition, it has been recently reported that pregnancy is associated with increased biological age, and that this increase is reversible postpartum^57,58^. Pregnancy is associated with reversible changes in blood cell composition, with changes in both total number and proportions of different cell types ^59–61^. In the analysis by Pham et al. (2024), adjusting the statistical models with estimated cell proportions attenuated the association between biological age and course of pregnancy. However, not all potentially relevant blood cell subtypes were accounted for in the analysis, and these findings should be replicated with measured, instead of estimated, blood cell proportions (see *Limitations and future perspectives*).

### Limitations and future perspectives

We show extreme and abundant differences for the values of ten BA indicators between the blood cell subtypes using four independent data sets. Importantly, we are able to show that the differences between the cell types appear to persist during adulthood, except for DunedinPACE. These results, together with the knowledge on wide ranges and age-associated changes in cell subtype proportions at population-level (Figure 4), highlight the need for additional efforts when using the existing epigenetic clocks or building up new ones. The cell composition in the blood samples may be accounted for in the statistical analysis if the composition is measured, however, measured cell type proportions are rarely available in large human cohort studies. One solution is to estimate the cell counts in a tissue sample using DNA methylation reference libraries for the various cell subtypes^62–64^. However, this cell count estimation is limited in two ways. First, DNA methylation-based cell count estimates may show only modest correlations with the cell counts obtained using other DNA methylation-based estimation algorithm^65^, and the reliability of the cell count estimation algorithms should be further evaluated in relation to e.g. FACS-based cell counts in larger, independent population cohorts. Second, current libraries do not cover all the different blood cell subtypes with diverse functionalities such as the more specific CD4+ ^66^ including regulatory T cell subpopulations^67^, or various B cell ^68^ or NK cell^69^ subpopulations. For example, NK cell subtypes show drastic changes in their abundance and/or functionality/properties in aging and/or age-related pathologies^70^. This limitation also extends to our analysis. Even though our observations are from sets of purified cell types that are often considered as ‘detailed cell separation’ (Figure 4), many potentially relevant blood cell subtypes couldn’t be analysed in our study because DNA methylation data is not available for them. Overall, our results highlight the need for analyses on the BA indicators in single cells.

In addition, even when the cell separation protocols and purity levels are according to the high standards in the field, cell subsets are hardly ever completely purified. In the four data sets used in this study, cells were separated using varying FACS protocols, and, for example, sometimes a cell subtype was determined with only one surface antigen while it was sometimes determined using more than one (Supplementary Table S1). The impurity may have influenced our results and caused noise in the cell subtype-specific BA values. Consistency in our findings suggest the extent of this noise is likely small but further studies are needed.

## Conclusions

Different blood cell subtypes generally show distinct biological ages (BAs), according to six BA indicators representing various aspects of biological aging. The differences between the cells can be substantial and they appear to persist across adult ages from 20 to 80 years for all BA indicators, except for DunedinPACE. When studying DNA methylation-based BA indicators in whole blood samples, the contribution of differing blood cell proportions needs to be considered. This is relevant for studies on physiological and pathological conditions known to have a significant effect on blood cell proportions, but especially for any potential aging interventions.

## Ethical statement

All datasets are in compliance with the Declaration of Helsinki and have been approved by local ethical committees, details can be found from the original publications (GSE35069 ^34^, GSE131989 ^35^, GSE166844 ^23^, GSE78942 ^36^ and SATSA ^37^.

## Author contributions

Conceptualization: SM, LKa; Methodology: SM, LKa; Formal analysis and investigation: SM, SR, JC, JM, LKa; Writing - original draft preparation: SM, LKa; Writing - review and editing: SM, SR, JC, JM, ISJ, LKu, SH, ER, LKa; Funding acquisition: SM, ER, LKa; Resources: SM, SH, ER, LKa; Supervision: SM, LKa.

## Supplementary material

Supplementary_Tables_S1-S8_2024-05-03.xlsx Supplementary_Figures_and_Results_2024-05-03.docx

