## Supplementary Figure for "Biological aging of different blood cell types"

- **Supplementary Figures S1-S3**
- **Supplementary Results**

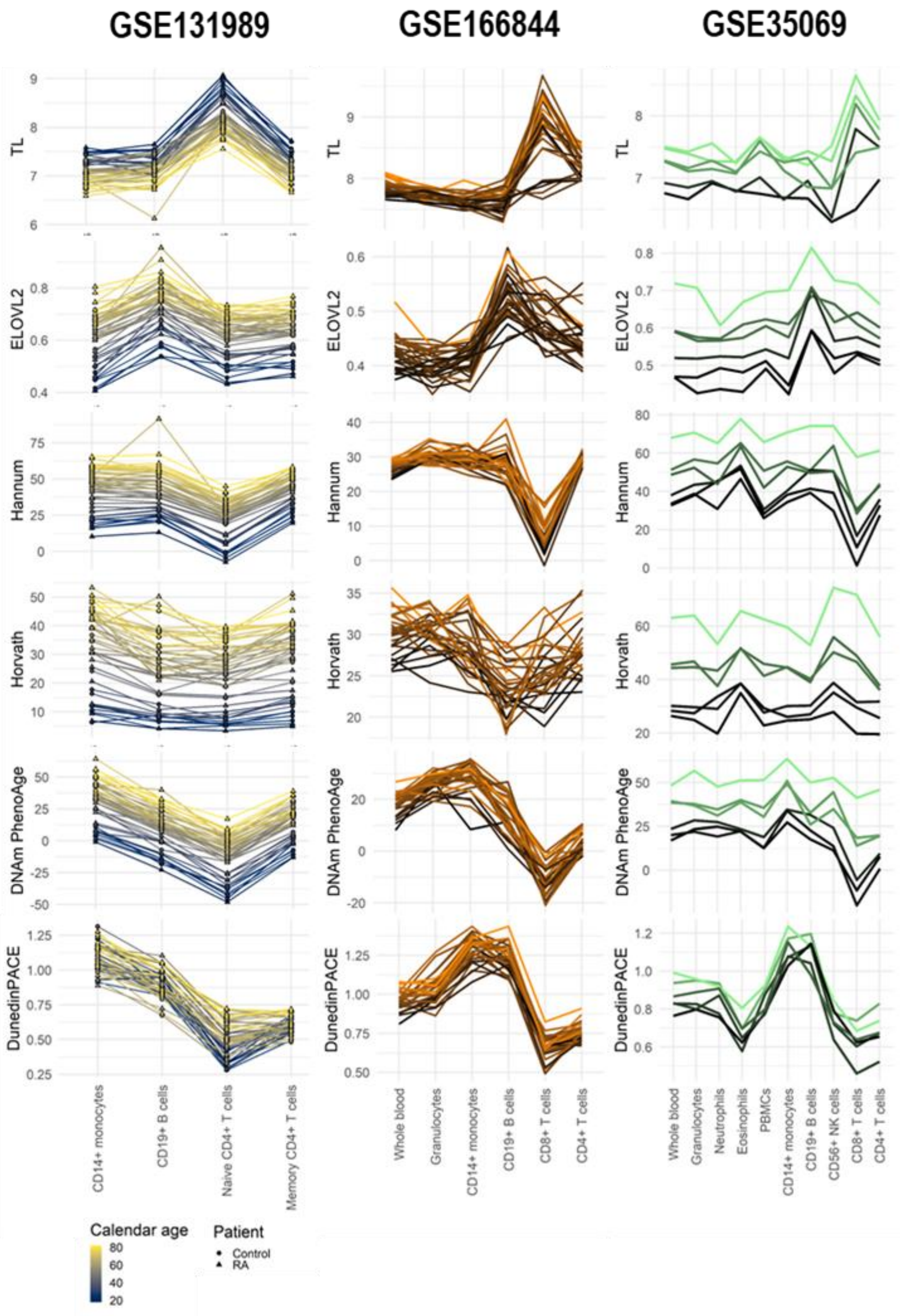


**Figure S1. Blood cell type-specific biological ages (BAs) and a BA rate**

DNA methylation-based indicators were assessed in three DNA methylation data sets (GSE131989, GSE166844, GSE35069) with 424 biological samples from 83 individuals and including 12 cell sample types. Lines are colored according to calendar age for GSE131989, and according to BA indicator values in whole blood sample for GSE166844 and GSE35069 because calendar age was either constant or not available. Each line represents one individual.

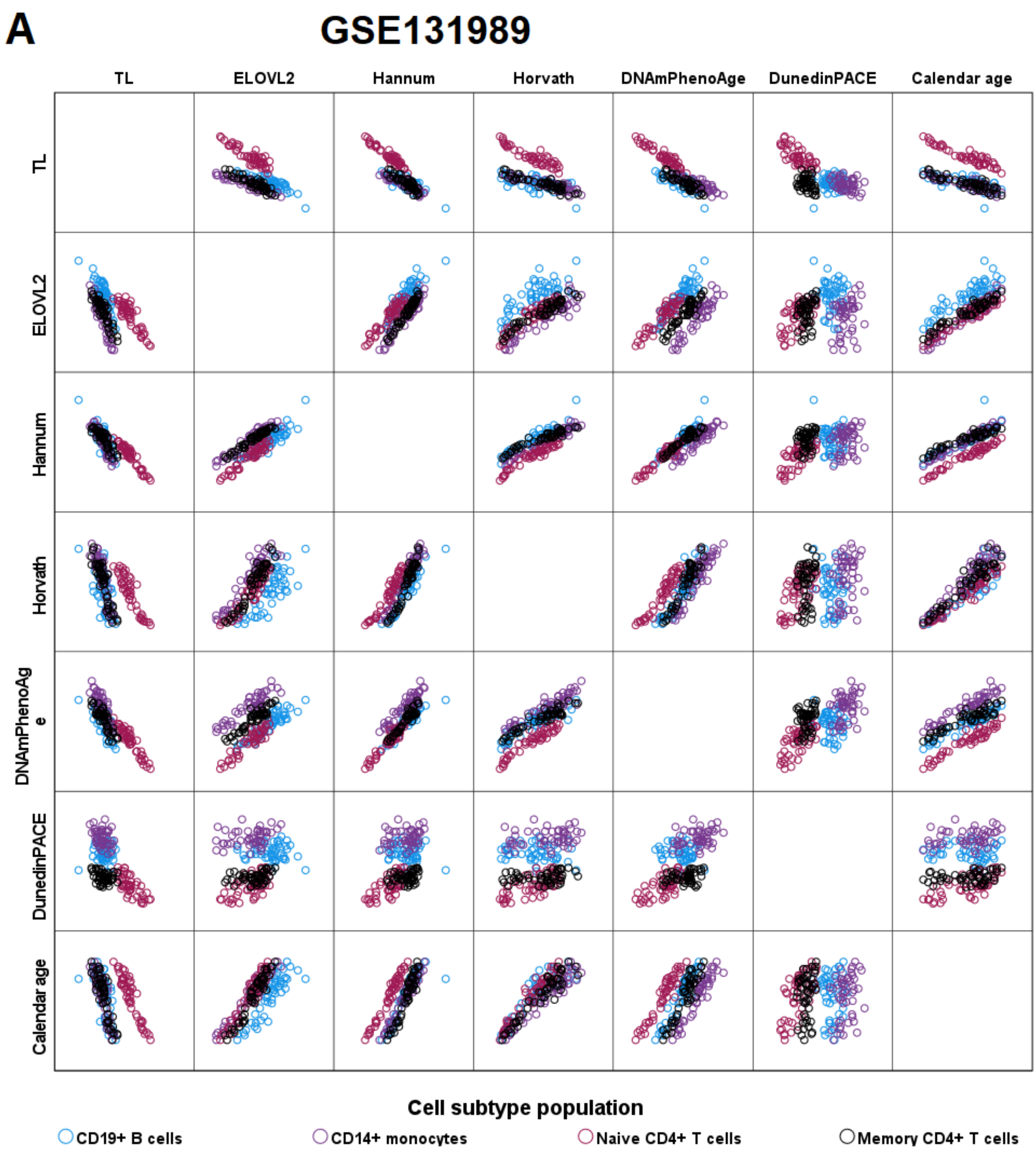

A
GSE131989
TL
ELOVL2
Hannum
Horvath
DNAmPhenoAge
DunedinPACE
Calendar age
TL
ELOVL2
Hannum
Horvath
DNAmPhenoAg
e
DunedinPACE
Calendar age
Cell subtype population
CD19+ B cells
CD14+ monocytes
Naive CD4+ T cells
Memory CD4+ T cells

## B

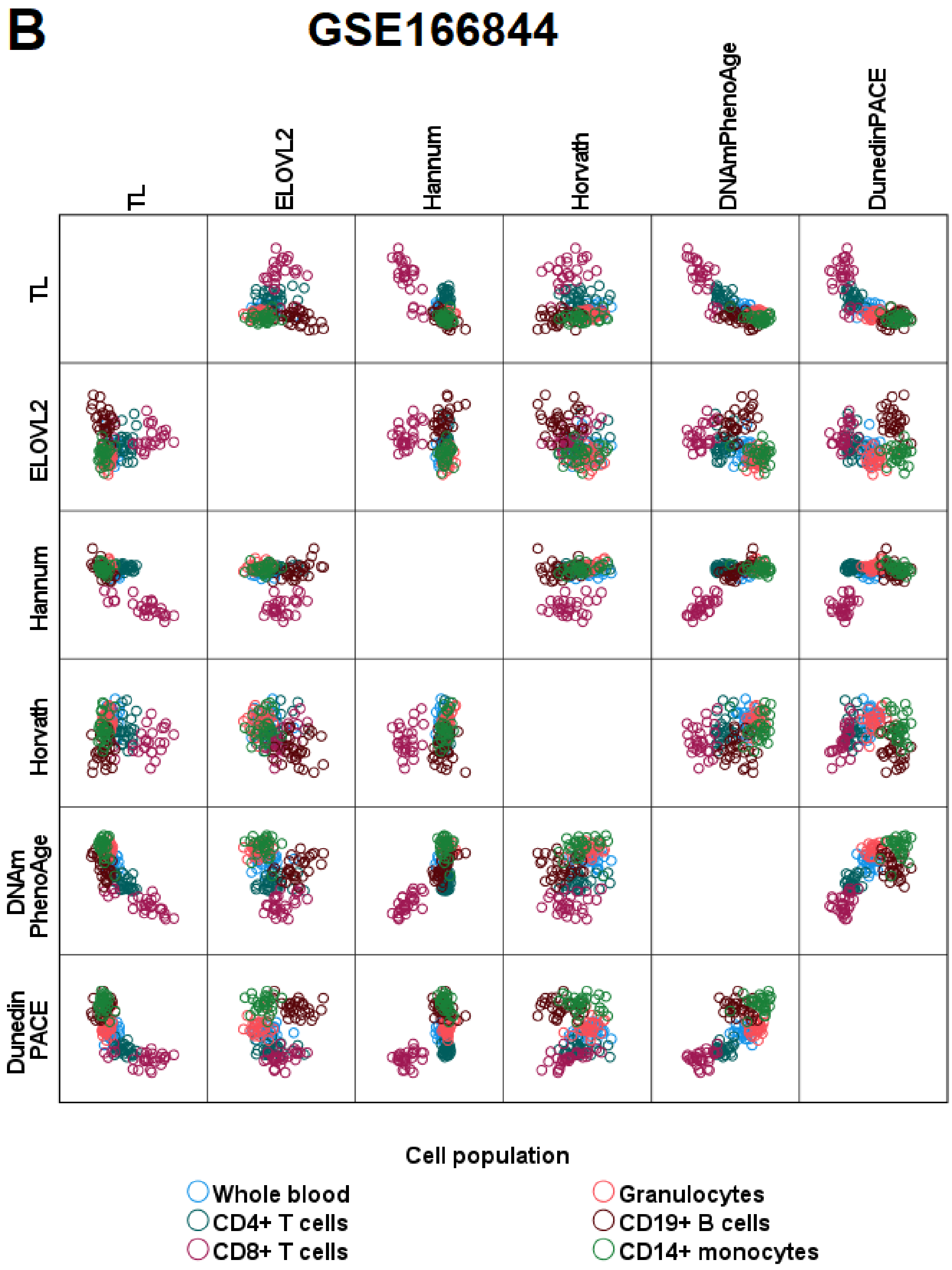

GSE166844
TL
ELOVL2
Hannum
Horvath
DNAmPhenoAge
DunedinPACE
DNAm PhenoAge
Dunedin PACE
Cell population
Whole blood
CD4+ T cells
CD8+ T cells
Granulocytes
CD19+ B cells
CD14+ monocytes

C

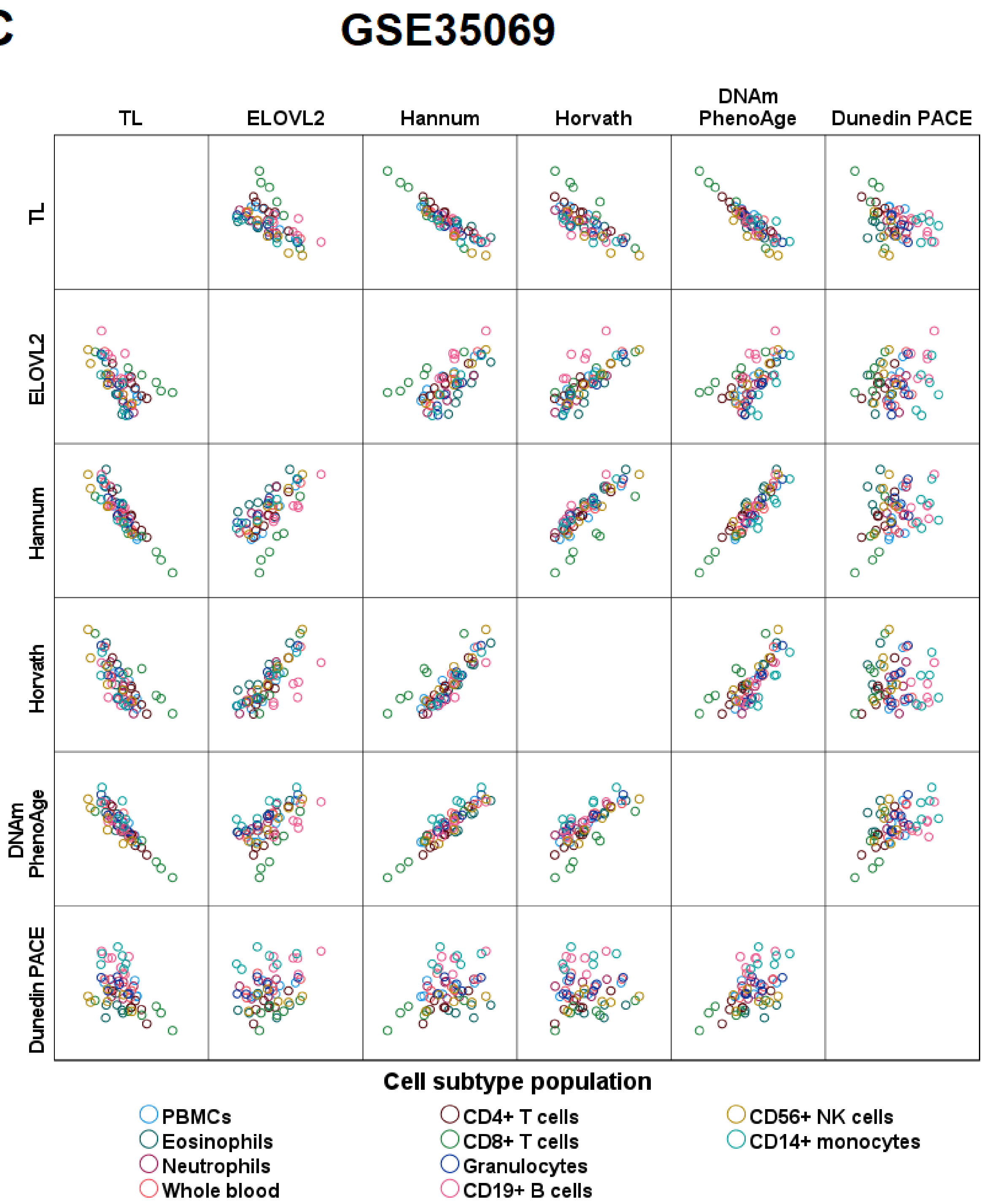


**Supplementary Figure S2. Relationship between the absolute values of the six BA indicators and calendar age in the three data sets, GSE131989 (A), GSE166844 (B) and GSE35069 (C)**

In each panel A-C, pairwise relationships in each data set are visualized as scatterplots. Thus, in panel A, there are 196, in panel B, 168 and in panel C, 60 biological samples which are colored according to cell subtype population. Correlation coefficients within each cell subtype population are shown in Supplementary Table S6-S8.

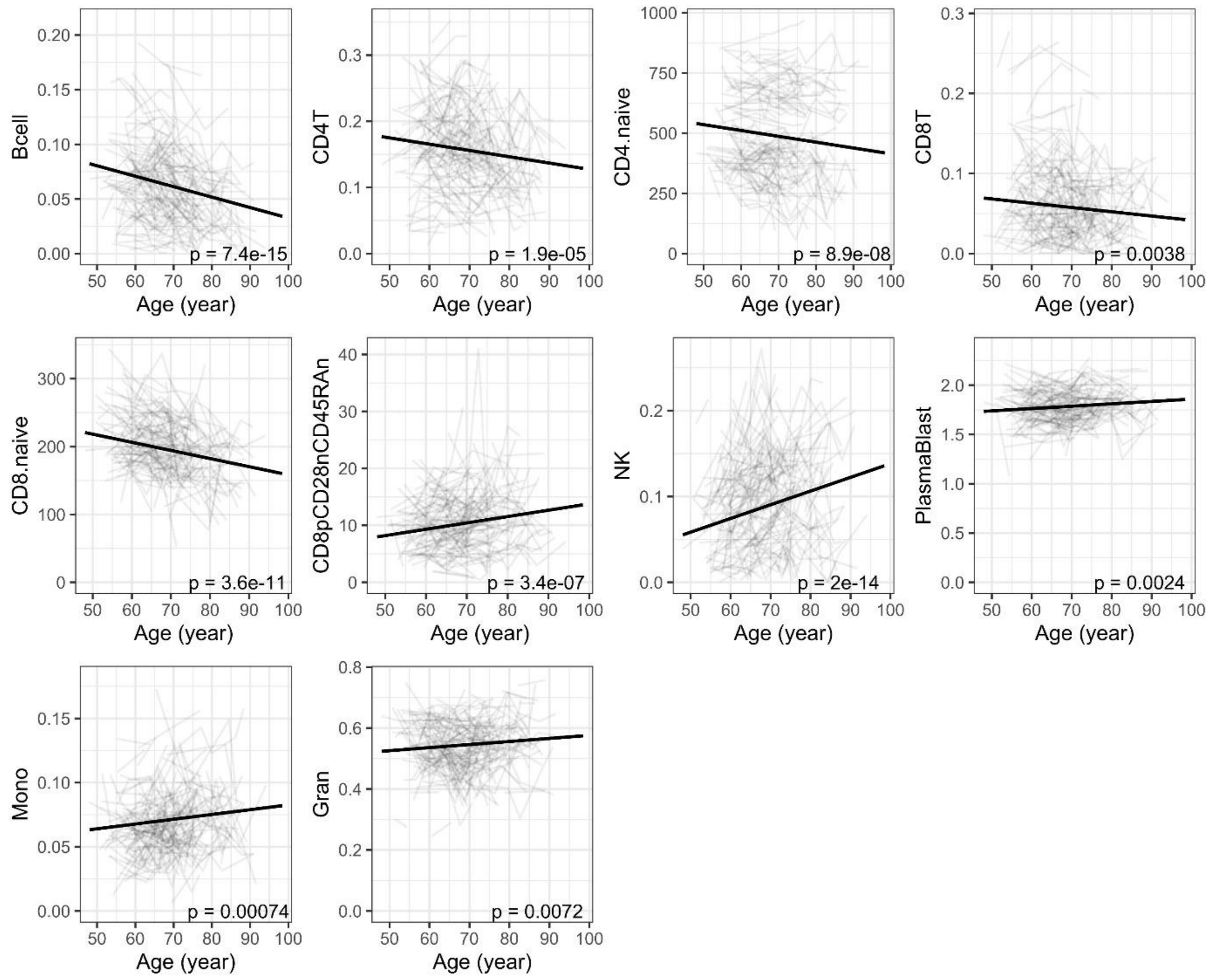


**Supplementary Figure S3. Cell count trajectories in blood with advancing calendar age**

Data from a longitudinal cohort (SATSA, n=328). p-value for yearly change from linear mixed model analysis. Cell counts were estimated from DNA methylation data (Wang et al., 2018). For this analysis, outlier values were filtered out from the cell count data rigorously (excluding values 0).

### Supplementary results

*Detailed description of the pairwise comparisons between cell types and whole blood*

*CD19+ B cells*

Methylation data in CD19+ B cells was available in three datasets. As compared to other cell types, CD19+ B cells displayed a statistically significant difference (Mann-Whitney U test p-value <0.05) in majority of the pairwise comparisons in GSE131989 and GSE166844 (Figure 2, Supplementary S3-S4). In the smallest dataset, GSE35069, statistically significant differences were mainly observed for DunedinPACE (Figure 2, Supplementary Figure S1, Supplementary Table S5). In summary, our results suggest CD19+ B cells are, according to the studied BA indicators, 'younger' as compared to CD14+ cells, but 'older' as compared to naïve CD4+ cells and total CD8+ T cells, although there are some discrepancies between the different BAs (Figure 2, Supplementary Table S3-S5). In comparison to whole blood, no clear pattern was observed for CD19+ B cells.

When comparing CD19+ B cells with CD14+ monocytes, lower values of Horvath, DNAmPhenoAge and DunedinPACE were detected in GSE131989 and GSE166844, suggesting that CD19+ cells are 'younger' as compared to CD14+ monocytes. In contrast, *ELOVL2* methylation was higher in CD19+ B cells as compared to CD14+ monocytes in GSE35069 and GSE131989, suggesting they are 'older', as *ELOVL2* hypermethylation has been shown to be associated with ageing(Garagnani et al., 2012) (Figure 2, Supplementary Tables S3-S5).

When comparing CD19+ B cells with CD4+ naïve T cells, we could observe a concordant pattern across different BA indicators suggesting CD19+ B cells are 'older' as compared to CD4+ naïve T cells. CD19+ B cells  had higher values of DNAmHannum, DNAmPhenoAge, DunedinPACE and *ELOVL2* methylation and lower value in DNAmTL as compared to CD4+ naïve T cells. When comparing CD19+ B cells to CD4+ memory T cells, a similar pattern of 'older' values for CD19+ B cells can be observed for DunedinPACE and *ELOVL2* methylation, both of which are higher in CD19+ B cells as compared to CD4+ memory cells.  In contrast, CD19+ B cells showed 'younger', i.e. lower value of Horvath as compared to CD4+ memory T cells (Figure 2, Supplementary Table S3).

When comparing CD19+ B cells with total CD4+ T cells, four out of the six BA indicators suggested CD19+ cells to be 'older' as compared to total CD4+ T cells, whereas two of the six

BA indicators suggested CD19+ B cells to be ‘younger’ as compared to total CD4+ T cells. ‘Older’ values for CD19+ B cells as compared to total CD4+ T cells were suggested by higher values of DunedinPACE and *ELOVL2* methylation and lower values of DNAmTL two datasets (GSE35069 and GSE166844) and by higher values of DNAmPhenoAge in one dataset (GSE166844). In contrast, ‘younger’ values for CD19+ B cells as compared to total CD4+ T cells were suggested by lower values of Horvath and Hannum (Figure 2, Supplementary Tables S4, S5).

When comparing CD19+ B cells with total CD8+ T cells, we could observe a clear pattern across different BA indicators suggesting CD19+ B cells are ‘older’ as compared to total CD8+ T cells. CD19+ B cells had higher values of Hannum, DNAmPhenoAge, DunedinPACE and *ELOVL2* methylation and lower value of DNAmTL as compared to total CD8+ T cells in GSE166844, and this was replicated for Hannum and DunedinPACE in GSE35069. The only contrasting BA indicator was Horvath, for which CD19+ B cells showed lower value in GSE166844 as compared to total CD8+ T cells (Figure 2, Supplementary Tables S4, S5).

When comparing CD19+ B cells with whole blood samples, there was no clear pattern regarding the direction of difference. Two of the BA indicators suggested CD19+ B cells to be ‘younger’ as compared to whole blood, as values of Horvath and DNAmPhenoAge were lower in CD19+ B cells as compared to whole blood in GSE166844. However, three of the BA indicators suggested CD19+ B cells to be ‘older’ as compared to whole blood, as values of DunedinPACE and *ELOVL2* methylation were higher in GSE166844 and GSE35069 and DNAmTL was lower in GSE166844 in CD19+ B cells as compared to whole blood (Figure 2, Supplementary Tables S4, S5).

*T cell subsets*

Data on various subsets of T cells was available in four datasets, including total CD4+ and CD8+ T cells (GSE166844 and GSE35069), CD4+ naïve and memory T cells (GSE131989) and CD4+CD28- and CD4+CD28+ T cells (GSE78942). Majority of pairwise comparisons across these cell types were statistically significant (Figure 2, Table 2, Supplementary Tables S3-S5). Our results suggest that CD8+ T cells are ‘younger’ as compared to CD4+ T cells, and that naïve CD4+ T cells are ‘younger’ as compared to memory CD4+ T cells (Figure 2, Supplementary Table S3-S5). In addition, CD4+CD28+ cells were identified to be ‘younger’ as compared to CD4+CD28- according to both BA indicators available for this dataset, Horvath and DNAmPhenoAge (Table 2), although for this data no statistical tests could be performed, as there were only four biological samples. As compared to whole blood, both CD4+ and CD8+ T cells are ‘younger’, although there are discrepancies between different BA indicators (Figure 2, Supplementary Table S3-S5). The magnitude of difference was larger between CD8+ T cells and whole blood as compared to CD4+ T cells and whole blood (Figure 1, Figure 2).

When comparing CD8+ T cells to CD4+ T cells, a clear pattern could be observed, suggesting that CD8+ T cells are ‘younger’ as compared to CD4+ T cells. Values for Horvath, Hannum, DNAmPhenoAge and DunedinPACE were lower and values for DNAmTL were higher in CD8+ T cells as compared to CD4+ T cells (GSE166844, Figure 2, Supplementary Table S4). In GSE35069 the difference between CD8+ and CD4+ T cells was not statistically significant, but a similar trend could be observed for Hannum and DNAmPhenoAge (Figure 2). In contrast, *ELOVL2* methylation values suggested CD8+ T cells to be ‘older’ as compared to CD4+ T cells, as the value of this BA indicator was higher for CD8+ T cells (GSE166844, Figure 2, Supplementary Table S4).

When comparing CD4+ naïve T cells to CD4+ memory T cells, we observed a concordant pattern suggesting that CD4+ naïve T cells are ‘younger’ as compared to CD4+ memory T cells. Values of Horvath, Hannum, DNAmPhenoAge and DunedinPACE were lower and DNAmTL were higher in CD4+ naïve T cells as compared to CD4+ memory T cells (GSE131989, Figure 2, Supplementary Table S3). Another CD4+ T cell subset we had data available were CD4+CD28+ and CD4+CD28- cells (GSE78942). As this dataset consisted of only 4 biological samples, no statistical test could be performed. However, CD4+CD28+ cells were identified to be ‘younger’ as compared to CD4+CD28- according to both BA indicators, Horvath and DNAmPhenoAge, available in that dataset (Table 2).

When comparing the different T cell populations to whole blood, we observed a clear pattern suggesting that CD8+ T cells are ‘younger’ as compared to whole blood sample. Values of Horvath, Hannum, DNAmPhenoAge and DunedinPACE were lower and value for DNAmTL was higher for CD8+ T cells as compared to whole blood (GSE166844, Figure 2, Supplementary Table S4). A similar pattern could be observed for Hannum and DunedinPACE in the small dataset GSE35069 (Figure 2, Supplementary Table S5). However, values for *ELOVL2* methylation in GSE166844 suggested an opposing pattern of CD8+ T cells being ‘older’ as compared to whole blood, as the value of this indicator was higher in CD8+ T cells (Figure 2, Supplementary Table S4).

When comparing CD4+ T cells to whole blood, results were similar to those for CD8+ T cells. Based on four of the six BA indicators, CD4+ T cells were suggested to be 'younger' as compared to whole blood. Values of Horvath, DNAmPhenoAge and DunedinPACE were lower and value of DNAmTL was higher for CD4+ T cells as compared to whole blood (GSE166844, Figure 2, Supplementary Table S4). A similar pattern was observed for DunedinPACE in GSE35069 (Figure 2, Supplementary Table S5). For CD4+ T cells, two BA indicators suggested the to be 'older' as compared to whole blood, as values for Hannum and *ELOVL2* methylation were higher for CD8+ T cells as compared to whole blood (Figure 2, Supplementary Table S4). For the majority of these differences, the magnitude of difference was larger between CD8+ T cells and whole blood as compared to CD4+ T cells and whole blood (Figure 1, Figure 2).

#### *CD14+ monocytes*

Data on CD14+ monocytes was available in three datasets. As compared to other cell types, majority of pairwise comparisons between CD14+ monocytes were statistically significant in GSE166844 and GSE131989 (Figure 2, Supplementary Figure S1, Supplementary Table S3 and S4). The prominent differences between CD14+ monocytes and CD19+ B cells have been described in detail in previous paragraph. Our results suggest that CD14+ monocytes are 'older' as compared to various T cell subsets, and also 'older' as compared to whole blood samples (Figure 2, Supplementary Tables S3-S5).

When comparing CD14+ monocytes to CD8+ T cells, we could observe a clear pattern suggesting that CD14+ monocytes are 'older' as compared to CD8+ T cells. Values for Horvath, Hannum, DNAmPhenoAge and DunedinPACE were higher and value for DNAmTL was lower for CD14+ monocytes as compared to CD8+ T cells in GSE166844, and the same pattern was observed for Hannum, DNAmPhenoAge and DunedinPACE in GSE35069 (Figure 2, Supplementary Tables S4, S5). In contrast, lower value of *ELOVL2* methylation suggested CD14+ monocytes to be 'younger' as compared to CD8+ T cells (GSE166844, Figure 2, Supplementary Table S4).

When comparing CD14+ monocytes to CD4+ T cells, three of the studied BA indicators suggested CD14+ monocytes to be older. Values of DNAmPhenoAge and DunedinPACE were higher and values of DNAmTL were lower in CD14+ monocytes as compared to CD4+ T cells in two datasets (GSE166844 and GSE35069, Figure 2, Supplementary Tables S4, S5). Only the lower value of *ELOVL2* methylation suggested CD14+ monocytes to be 'younger' as compared to CD4+ T cells (GSE166844, Figure 2, Supplementary Table S4).

When comparing CD14+ monocytes to naïve and memory subsets of CD4+ T cells, a similar pattern as in total CD4+ T cells can be observed, that is that CD14+ are 'older'. Values of Horvath, Hannum, DNAmPhenoAge and DunedinPACE are higher and value of DNAmTL is lower in CD14+ monocytes as compared to CD4+ naïve T cells. Values of DNAmPhenoAge and DunedinPACE are higher in CD14+ monocytes as compared to CD4+ memory T cells (GSE131989, Figure 2, Supplementary Table S3).

When comparing CD14+ monocytes to whole blood, we observed a concordant pattern suggesting CD14+ monocytes are 'older' as compared to whole blood sample. Values of Hannum, DNAmPhenoAge and DunedinPACE were higher and value of DNAmTL was lower in CD14+ monocytes as compared to whole blood (GSE166844, Figure 2, Supplementary Table S4). For DunedinPACE, this finding was replicated in GSE35069 (Figure 2, Supplementary Table S5).
